# Phylogenetic incongruence and the origins of cardenolide-resistant forms of Na^+^, K^+^- ATPase in North American *Danaus* butterflies

**DOI:** 10.1101/048033

**Authors:** Matthew L. Aardema, Peter Andolfatto

**Affiliations:** Department of Ecology and Evolutionary Biology, Princeton University, Princeton, NJ, 08544, USA; Current Address: American Museum of Natural History, New York, NY, 10024, USA; Lewis-Sigler Institute for Integrative Genomics, Princeton University, Princeton, NJ, 08544, USA

**Keywords:** *Danaus*, Danaidae, milkweed butterfly, incomplete lineage sorting, genetic introgression, Na^+^, K^+^-ATPase

## Abstract

Rapid species radiations can obscure phylogenetic relationships between even distantly related species and lead to incorrect evolutionary inferences. For this reason, we examined evolutionary relationships among the three North American milkweed butterflies, *Danaus plexippus, D. gilippus* and *D. eresimus* using >400 orthologous gene sequences assembled from transcriptome data. Contrary to previous phylogenetic assessments, our results indicate that *D. plexippus* and *D. eresimus* are the sister taxa among these species. This result explains many previously noted phylogenetic incongruences such as an amino acid substitution in the sodium-potassium pump (Na^+^,K^+^-ATPase) of *D. eresimus* and *D. plexippus*, which increases resistance to the toxins found in these butterflies’ host plants. In accordance with a rapid radiation of *Danaus* butterflies, we also find evidence that both incomplete lineage sorting and post-speciation genetic exchange have contributed significantly to the evolutionary histories of these species. Furthermore, our findings suggest that *D. plexippus* is highly derived both morphologically and behaviorally.

## Introduction

Plants in the genus *Asclepias* (milkweed) often produce a class of toxic, secondary metabolites called cardenolides, which provide defense against herbivory (Dobler et al. 2011; Agrawal et al. 2012). Despite these protective chemicals, many insect species have evolved the ability to feed on these plants. The phylogenetically broad insect herbivore community that specializes on *Asclepias* has proved to be a particularly good system for studying convergent evolution (Dobler et al. 2012; Zhen et al. 2012; Petschenka et al. 2013). Studies of convergence in these insects have improved our understanding of the underlying genetic architecture behind adaptive traits and given insights into the extent of constraint on genetic change (Stern 2013; Dobler et al. 2015).

As larvae, butterflies of the genus *Danaus* feed almost exclusively on *Asclepias* host plants (Ackery and Vane-Wright 1984). Many species also sequester the plants’ toxins for their own defense (Brower and Moffit 1974; Ackery and Vane-Wright 1984). Within the three species of North American *Danaus*, both *D. plexippus* (the monarch) and *D. eresimus* (the soldier) appear to be fixed for a functionally important amino acid substitution in the H1-H2 extracellular domain of Na^+^K^+^‐ ATPase (N122H; Fig. S1; Zhen et al. 2012). This substitution decreases larval sensitivity to cardenolides consumed while feeding on *Asclepias* host plants, and may also facilitate toxin sequestration (Dobler et al. 2012; Zhen et al. 2012; Petschenka et al. 2013). The N122H substitution has not been observed in the third North American *Danaus, D.gilippus* (the queen; Holzinger and Wink 1996; Aardema et al. 2012; Dobler et al. 2012; Zhen et al. 2012; Petschenka et al. 2013).

Interestingly, among these three species, *D. gilippus* and *D. eresimus* have historically been considered sister taxa, while *D. plexippus* has been classified as a more distant relative within the *Danaus* genus (Fig. S1; Ackery and Vane-Wright 1984). This relationship was initially based on a large number of shared morphological traits and behavioral similarities between *D. gilippus* and *D. eresimus*, and it received further supported in analyses using sequence data from a small number of mitochondrial (mtDNA), ribosomal (rRNA) and nuclear genes (Lushai et al. 2003; Smith et al. 2005; Brower et al. 2010). However, in addition to the N122H substitution shared by *D. eresimus* and *D. plexippus*, other apparent phylogenetic incongruences have also been noted. For example, *D. eresimus* and *D. plexippus* are each reported to have 30 chromosomes, whereas *D. gilippus* appears to typically have 29 (Brown et al. 2004). Also, *D. eresimus* and *D. plexippus* have a shared Malic enzyme (ME) allozyme phenotype distinct from *D. gilippus* (Kitching 1985). A third example of apparent phylogenetic incongruence in these butterflies are the rear projections on the eighth sternal segment of *D. eresimus* and *D. plexippus* males, which *D. gilippus* males lack (Ackery and Vane-Wright 1984).

Given the inferred species phylogeny that places *D. gilippus* and *D. eresimus* as closer relatives within *Danaus*, the origins of the N122H substitution and other incongruences are unclear. In the case of N122H, it is possible that this mutation arose independently and in parallel within both *D. plexippus* and *D. eresimus*, and then subsequently fixed in each butterfly. Examples of this specific mutation evolving in disparate taxa have been reported previously, in other insect taxa, although not in other Lepidoptera outside the *Danaus* genus (Aardema et al. 2012; Zhen et al. 2012). It is also possible that both the asparagine (N) and histidine (H) alleles were segregating in the common ancestor of these three species. If this is the case, then the histidine must have fixed independently in *D. plexippus* and *D. eresimus*, while being lost in *D. gilippus*. Such incomplete lineage sorting (ILS) is a common cause of phylogenetic incongruence (Degnan and Rosenberg 2006; 2009; Maddison and Knowles 2006). Adaptive introgression is a third hypothesis to explain fixation of the N122H allele in both *D. plexippus* and *D. eresimus*. Genetic exchange after splitting has become increasingly recognized as an important contributor to the evolutionary history of species (reviewed in Mallet et al. 2015), and it can often lead to phylogenetic incongruence (e.g. Bachtrog et al. 2006; Baack and Rieseberg 2007; Cui et al. 2013).

It is also possible that while the many morphological and behavioral similarities of *D. eresimus* and *D. gilippus* suggest they are more closely related, our understanding of the phylogenetic relationships of North American *Danaus* may benefit from a fresh examination. For this reason, the first goal of this study was to re-evaluate species relationships among these butterflies using a much larger set of genetic sequences than has previously been employed. We also compared the extent to which ILS and genetic introgression may have contributed to phylogenetic incongruences between these species. Both ILS and genetic introgression have previously been proposed to explain observed morphological and genetic incongruences in the *Danaus* genus (Lushai et al. 2003; Smith et al. 2005), but to date neither hypothesis has been examined explicitly.

Our results indicate that in contrast to previous phylogenetic analyses, *D. eresimus* and *D. plexippus* are the sister taxa among North American *Danaus* butterflies. This suggests that the N122H mutation in both species can be explained by an origin in their shared, common ancestor. However, we also find that levels of genetic incongruence in these species are high, and may result from both ILS and post-splitting genetic exchange. In addition to the origins if the N122H mutation in *Danaus*, the findings of this study indicate that *D. plexippus* is highly derived morphologically and behaviorally, relative to other *Danaus* species.

## Materials and Methods

### Data preparation and location of orthologous sequences

To examine phylogenetic incongruences and the origins of the N122H substitution within *Danaus*, we took advantage of previously produced RNAseq data for the three *Danaus* species present in North America and a fourth milkweed butterfly outgroup, *Lycorea halia* (Zhen et al. 2012). Using just three focal species allows us to compare a major topology to only two possible minor topologies in a series of four-taxa trees, and assess whether these minor topologies occur at similar or different frequencies to one another. Such an assessment is a common way to investigate the origins of phylogenetic incongruence (e.g. Green et al. 2010; Eriksson and Manica 2012).

### Reference-mapped alignments

Raw RNAseq reads from each of the four species were trimmed for quality (Phred QV ≥ 20) and contiguous length (≥30 nucleotides). We first mapped the trimmed reads from each sample to *D. plexippus* reference coding sequences (CDS; http://monarchbase.umassmed.edu/resource.html) using Stampy (v. 1.0.17; Lunter and Goodson 2011) with default parameters except substitution rate, which we set to 0.01. We used SAMtools (v. 0.1.4) to convert the resulting SAM formatted files to BAM formatted files while filtering for mapping quality (MAPQ > 20, Li et al. 2009). The MarkDuplicates utility in Picard tools (v. 1.77; http://broadinstitute.github.io/picard/) was used to remove PCR duplicates. We used the HaplotypeCaller utility in GATK (v. 3.3; McKenna et al. 2010) to call variants, and then filtered all sites for a minimum read depth of 30. All sequences were checked by eye to ensure indels were in frame with relation to the reference.

### De novo-assembled alignments

For a second set of sequence alignments, we produced independent *de novo* transcriptome assemblies for each species using the programs Velvet (v. 1.2.10) and Oases (v. 0.2.08) with a kmer length of 31 and a minimum read depth of 10 (Zerbino and Birney 2008; Schulz et al. 2012). In cases where multiple isoforms were assembled, we retained only the longest one. We located mtDNA regions by blasting each of the four species’ transcripts against the *D. plexippus* mitochondrial genome (Genbank accession number KC836923; ‘blastn’; Altschul et al. 1990). This was done for comparison to previously published results (Lushai et al. 2003; Smith et al. 2005; Brower et al. 2010). To locate orthologous nuclear sequences from our assembled transcriptomes, the four transcript sets were compared to amino acid sequences of predicted proteins for *D. plexippus* (Dp_geneset_OGS2_pep.fasta, http://monarchbase.umassmed.edu/resource.html), using Blast with a translated nucleotide query (‘blastx’) and a minimum e value of 1e-50. We compared our transcripts to amino acid sequences to improve likely ortholog recognition in the outgroup, *L. halia*. Only Blast hits that were at least 100 nucleotides long were retained for subsequent ortholog comparison between the four species. This was done to reduce regions that had limited phylogenetic information in our dataset, as well as improve the accuracy of our alignments (Talavera and Castresana 2007). For regions where any species had more than one transcript matching the *D. plexippus* reference protein sequence, we discarded this region for all species to avoid potential gene duplicates. Generally, a low level of polymorphism within a sample does not affect contig assembly in Velvet, and in these cases only a single allele is retained (Zerbino and Birney 2008). However, it is possible that highly polymorphic regions will assemble into more than one distinct transcript. In our pipeline, such regions would resemble gene duplicates and subsequently be removed.

We aligned orthologous regions based on amino acid similarity with MUSCLE (Edgar 2004) as implemented in SeaView (v. 4.5.4; Gouy et al. 2010). We then checked each sequence by eye and manually trimmed them to the length of the shortest region observed among the four species. As we are examining the relationship between three focal species and an outgroup, a missing species for a genetic region would have limited utility for phylogenetic inferences. We made no effort to identify specific genes within these datasets, with the exception of Na^+^,K^+^-ATPase which we located in our *de novo* assemblies for each species using Blast (see above).

### Species trees

We first concatenated all aligned mtDNA regions into a single dataset (i.e. no partitioning between regions). With jModelTest 2 (v. 2.1.7, Guindon and Gascuel 2003; Darriba et al. 2012), we determined that a the generalized time reversible model (GTR; Tavaré 1986), with a gamma distribution of rate heterogeneity best fit this data based on AIC score (i.e. GTR + Γ model). We also concatenated all loci into single alignments (i.e. no partitioning between genes) independently for both nuclear gene datasets (reference-mapped and *de novo*-assembled). As above, we used jModelTest 2 to determine the best-fitting mutation model. For both genomic datasets, the best fit was GTR model with a proportion of invariable sites and a gamma distribution of rate heterogeneity (i.e. GTR + I + Γ model). Finally, we examined the topology of Na^+^,K^+^-ATPase using our *de novo* assemblies of this gene for each species. In this case the GTR + Γ model best fit the data.

For all datasets, we independently produced maximum likelihood phylogenies in RAxML (v. 8.1.7, Stamatakis 2014) with 100 bootstrap replicates for each of the three possible topologies among the *Danaus* species. The site log likelihoods from RAxML were used to perform the approximately unbiased (AU) test implemented in CONSEL 0.2 (Shimodaira and Hasegawa 2001). Two trees were statistically different from one another if p ≤ 0.05.

### Discordance among nuclear gene trees

To quantify the extent of phylogenetic discordance among our nuclear genes, we employed Bayesian concordance analysis (BCA), as implemented in BUCKy (v. 1.4.3, Larget et al. 2010), to determine what proportion of genes concur with the primary species tree, and what proportions support the two alternative phylogenetic relationships (Ané et al. 2007). While BUCKy can identify significant levels of gene-tree discordance, it does not differentiate between ILS and hybridization to explain this discordance (Larget et al. 2010).

To carry out the BCA, we first produced individual trees for each gene in the dataset of *de novo*-assembled transcripts using MrBayes (v. 3.2.2, Ronquist and Huelsenbeck 2003), with two independent runs of 10^7^ generations and a “burn in” period of 10^5^ generations. Rather than *a priori* selecting a mutation model for each gene, we used the reversible-jump Markov chain Monte Carlo approach as described by Huelsenbeck and colleagues to examine the parameter space for each locus and find the best set of parameters for that particular dataset (Huelsenbeck et al. 2004).

We combined each of the two independent MrBayes runs for each gene using the mbsum command in BUCKy (v. 1.4.3, Larget et al. 2010). With the combined files, we examined the extent of discordance as measured by the sample-wide concordance factor (CF). This measure indicates the proportion of gene trees that support a particular species tree. The data was analyzed using an α prior of 1. α is an *a priori* discordance parameter that indicates the expected level of discordance among the different genes being analyzed (ranging from 0 to ∞; Larget et al. 2010). If there is no expected discordant topologies among gene trees, then α=0, whereas α=∞ indicates that all gene trees are expected to have independent topologies. In our analysis, other values of α (0.1 & 10) did not greatly alter inferred levels of incongruence (SI Table 3).

### Differentiating between incomplete lineage sorting and genetic introgression

While BUCKy analysis reveals the extent of nuclear discordance among taxa, it does not indicate whether observed discordance is more likely to be due to ILS or genetic introgression after species splitting (although these are not mutually exclusive). To examine these two hypotheses we employed the ABBA/BABA test to calculate Patterson’s D statistic for our dataset (Green et al. 2010). This statistic uses comparisons between three focal samples and an outgroup to determine whether phylogenetically-informative sites are in agreement with the primary phylogeny (AABB sites), or else support one of the two possible alternative relationships (ABBA or BABA, respectively). If incomplete lineage sorting is the primary contributor to observed phylogenetic incongruence, then ABBA and BABA sites should be present in approximately equal frequencies (D statistic ≈ 0). However, genetic exchange that occurs after splitting may result in an excess of either ABBA or BABA sites (D statistic ≠ 0). Structure in the ancestral population could also produce an asymmetry in ABBA/BABA sites, whereas post-splitting, symmetrical genetic exchange could produce equal frequencies of sites (Durand et al. 2011).

The ABBA/BABA test was performed with all informative sites, 4-fold synonymous sites and 0-fold replacement sites from our 478 *de novo*-assembled gene set. The standard deviation of D was determined by block jackknife sampling (Kunsch 1989) with a block size of 40 genes, giving us 12 independent block estimates of D. 95% confidence intervals based on this jackknifed dataset were used to assess if D differed significantly from 0.

### Genetic diversity and divergence times

ILS is most likely to be observed when multiple speciation events occur in rapid succession. Therefore, we wanted to estimate approximate splitting times for these species. However, changes in the effective population size of a species (N_e_) over time can influence divergence estimates and patterns of evolution (Charlesworth 2009). Therefore, we examined relative differences in contemporary N_e_ for each species by calculating 4-fold synonymous genetic diversity (π_4f_, Nei and Li 1979) within each of the four individual, wild-caught samples. To do this, we first mapped our trimmed reads to the species specific, *de novo*-assembled transcriptome for each of the four butterflies. We then used the HaplotypeCaller utility in the program GATK to identify genetic variants (McKenna et al. 2010). After this, sites with a read depth <30 were filtered out of the dataset. Likely coding sequences were determined by independently comparing each of the four species’ *de novo* assembled transcriptomes to the set of *D. plexippus* predicted proteins (Dp_geneset_OGS2_pep.fasta, http://monarchbase.umassmed.edu/resource.html), using Blast with a translated nucleotide query (‘blastx’; Altschul et al. 1990) and a minimum e value of 1e-50.

Using a custom Perl script, we calculated mean genetic diversity per gene for all 4-fold synonymous sites (π_4f_) using the formula: (n/n-1) 2p(1-p), where n is the number of chromosomes in the sample (here, n = 2) and p and (1-p) are the frequencies of alleles observed at a given site (here, either 1 & 0 [homozygous sites] or 0.5 & 0.5 [heterozygous site]). For each species, mean π_4f_ across all genes and 95% confidence intervals were estimated by bootstrap sampling each gene set 1,000 times with replacement, weighted by the number of 4-fold synonymous sites observed in each gene.

Species splitting times (t) can be roughly calculated using the formula t = dS – π/2μ, where dS is an estimate of the average number of synonymous nucleotide differences per synonymous site observed between two species (Nei 1987), π is a measure of intraspecific diversity, and μ is an estimate of the mutation rate (Hudson et al. 1987; Bachtrog et al. 2006). For each species pair we estimated dS for all synonymous sites with the method of Nei and Gojobori (1986) as implemented in PAML (v. 4.8; Yang 2007). For our estimate of π, we used the larger of the species’ π_4f_ estimates from each species pair (Table S4, following Bachtrog et al. 2006). No estimates of a mutation rate for *Danaus* butterflies have been made, so we used a recent, experimentally determined per generation estimate from *Heliconius melpomene* for our calculations (2.9 x 10^-9^ per site, Keightley et al. 2015). Both *Danaus* and *Heliconius* species are in the Nymphalidae family of butterflies. This per generation mutation estimate was converted to a per year estimate (μ = 1.45 x 10^-8^), conditioned on five generations per year (Malcolm et al. 1987).

We additionally employed a Bayesian method to examine approximate species’ splitting times. As there are no known fossils that would allow us to independently calibrate any of the nodes in our phylogeny, we employed a rate-based estimate of evolution at neutral sites following methods described by Obbard and colleagues (Obbard et al. 2012). For evaluation of neutral evolution, we utilized all 4-fold synonymous sites from the concatenated dataset of our *de novo* assemblies. While mutations at these sites may not be entirely neutral due to processes such as codon usage bias (Hershberg and Petrov 2008) and other types of selection (Lawrie et al. 2013), they are the best available option in this dataset.

Because we concatenated all 4-fold synonymous sites from each gene into a single dataset, we did not take into account variation in evolution rates between genes. Rather, we used the estimated 95% confidence intervals of the mutation rate given for *H. melpomene* to produce a lognormal distribution of rate variation around the mean for the complete dataset (6.5 x 10^-9^ – 2.8 x 10^-8^ changes per year, again conditioned on five generations per year [Malcolm et al. 1987]). This allowed us to partially capture potential variation in rates among branches and may reduce overestimating the length of shorter branches (Schwartz and Mueller 2010).

We ran two MCMC chains of 10^8^ iterations in the program BEAST (v. 1.8.1, Drummond and Rambaut 2007), with a log-normal relaxed molecular clock. Our substitution model was that of Hasegawa, Kishino and Yano (HKY, Hasegawa et al. 1985), with a proportion of invariant sites and a gamma distribution of rate heterogeneity. We used a starting tree that matched the one found in our previous analyses based on nuclear genes (Fig. 1B-D), with a birth-death process of lineage birth as our tree prior (Gernhard 2008). Our “burn in” period was the initial 10% of states, and parameters were logged every 1,000 iterations. LogCombiner was used to merge our two separate runs (v. 1.8.1, included with the BEAST package). The log files were checked using Tracer (v. 1.6; Rambaut et al. 2014) to ensure that an effective sampling size (ESS) greater than 200 were achieved for each parameter. Divergence times were estimated based on the 95% highest posterior density (HPD) interval.

**Figure 1.**
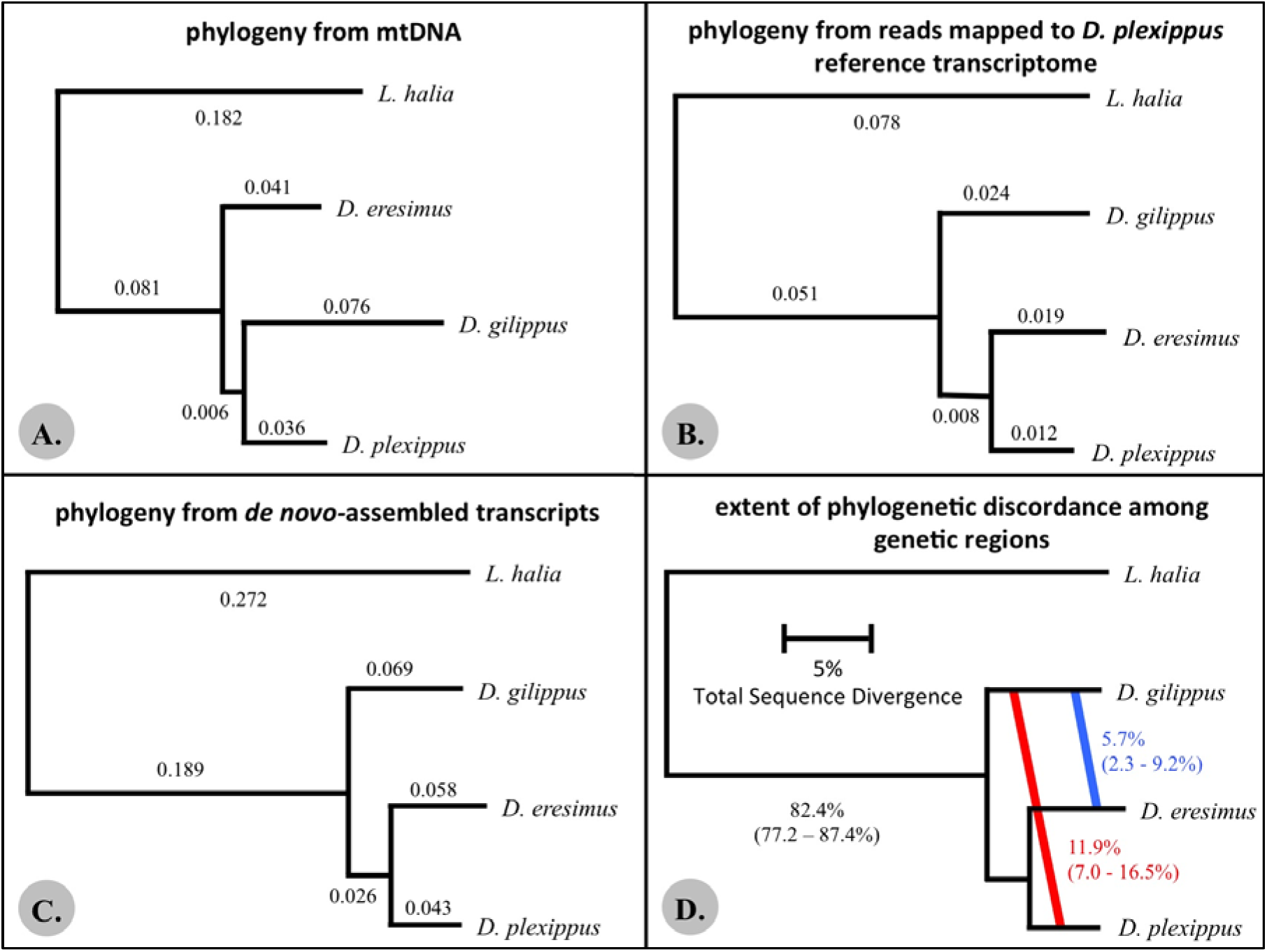
A) The best supported phylogenetic relationship as determined using mtDNA sequence data. Numbers indicate branch lengths. This tree topology had 65% bootstrap support. B) The best supported phylogenetic relationship produced from 471 reference-mapped nuclear gene sequences. Numbers indicate branch lengths. This tree topology had 100% bootstrap support. C) The best supported phylogenetic relationship as determined using data from 478 *de novo*-assembled nuclear gene sequences. Numbers indicate branch lengths. This tree topology had 100% bootstrap support. D) Levels of phylogenetic concordance/discordance between the best-supported species tree and individual gene trees. The red text and diagonal line indicates the level of discordance (with credibility intervals) that supports a closer relationship between *D. plexippus* and *D. gilippus* (with α = 1.0). The blue text and diagonal line indicated the level of discordance (with credibility intervals) that supports a closer relationship between *D. gilippus* and *D. eresimus* (with α = 1.0). **NOTE: Branch lengths in these trees (A-D) are not drawn to the same scale *between* trees.**

## Results

### Phylogenic analysis

Previous studies based on morphology and limited genetic data concluded that *D. eresimus* and *D. gilippus* are more closely related to one another than either is to *D. plexippus* (Ackery and Vane-Wright 1984; Lushai et al. 2003; Smith et al. 2005; Brower et al. 2010). Here, we re-evaluated this conclusion using multiple datasets of nuclear coding sequences, as well as mitochondrial data. The mitochondrial dataset totaled 6,648 aligned nucleotides. The reference-mapped, nuclear gene dataset contained 471 sequences, totaling 62,646 nucleotides, with an average sequence length of 133 nucleotides. The *de novo* assembled nuclear gene dataset contained 478 sequences, totaling 220,188 nucleotides, with an average sequence length of 461 nucleotides. All datasets were limited in the total number of genes analyzed as overlapping regions in each gene had to be assembled or mapped in all four species for inclusion.

In contrast to previous findings (Lushai et al. 2003; Smith et al. 2005, Brower et al. 2010), the best-supported mtDNA tree suggested that *D. plexippus* and *D. gilippus* are sister taxa and *D. eresimus* is basal to them (Fig. 1A). However, the log likelihoods of the three possible mtDNA trees were not significantly different from each other (Table 1) and the bootstrap support for this relationship was only 65%.

**Table 1.** Comparison of the three possible phylogenetic relationships among the butterflies in this study for all mitochondrial (mtDNA) and genomic regions (gDNA). [“ref. mapped” = reference-mapped nuclear gene dataset; *“de novo”* = *de novo-assembled* nuclear gene dataset]. Significance between trees was determined using the AU test. See Materials and Methods for more details.

| Dataset | Tree Comparison<br>(see Fig. 2) | $\vert \Delta \ln L\vert $ | p value from AU<br>test |
| --- | --- | --- | --- |
| mtDNA | Tree A – Tree B | 1.5 | 0.270 |
| mtDNA | Tree A – Tree C | 0.2 | 0.391 |
| mtDNA | Tree B – Tree C | 1.6 | 0.202 |
| gDNA (ref. mapped) | Tree A – Tree B | 267.4 | <0.001* |
| gDNA (ref. mapped) | Tree A – Tree C | 239.2 | <0.001* |
| gDNA (ref. mapped) | Tree B – Tree C | 30.3 | 0.024* |
| gDNA (*de novo*) | Tree A – Tree B | 282.9 | <0.001* |
| gDNA (*de novo*) | Tree A – Tree C | 297.6 | <0.001* |
| gDNA (*de novo*) | Tree B – Tree C | 22.6 | 0.057 |
| Na<sup>+</sup>,K<sup>+</sup>-ATPase | Tree A – Tree B | 3.5 | 0.841 |
| Na<sup>+</sup>,K<sup>+</sup>-ATPase | Tree A – Tree C | 3.5 | 0.220 |
| Na<sup>+</sup>,K<sup>+</sup>-ATPase | Tree B – Tree C | 4.4 | 0.069 |

\* indicates statistical significance at $p \leq 0.05$ .

For both the reference-mapped and *de novo*-assembled datasets, the best-supported topology placed *D. plexippus* and *D. eresimus* as sister taxa, with *D. gilippus* a more distant relative (Fig. 1B & C). For the reference-mapped dataset, a relationship that places *D. eresimus* and *D. plexippus* as sister taxa was significantly better than the other two possible topologies, whereas the relationship that places *D. plexippus* and *D. gilippus* as sister taxa was significantly better than the relationship that placed *D. gilippus* and *D. eresimus* as sister taxa (Table 1). For the *de novo* assembled dataset, the two alternative topologies were significantly worse fits than the primary tree, but they were not significantly different from each other (Table 1). There was 100% bootstrap support for the best tree topology in both analyses utilizing nuclear gene datasets.

The major difference between the trees produced by the two nuclear gene datasets (reference-mapped vs. *de novo*-assembled) is in the estimated branch lengths. Specifically, the branch lengths are substantially shorter in the reference-mapped dataset, especially for the ancestral *Danaus* branch and that of the outgroup, *L. halia*. We attribute these differences to a bias towards highly conserved genes in the reference-mapped dataset (Hornett and Wheat 2012). Such bias becomes more problematic as the evolutionary distance between two species increases. Additionally, for all four species, fewer reads mapped to the *D. plexippus* reference than to each species’ independently assembled *de novo* transcriptome (SI Table 2), and the average sequence length was shorter. For these reasons, in subsequent analyses, we exclusively used the data from our *de novo*-assembled dataset.

**Table 2.**
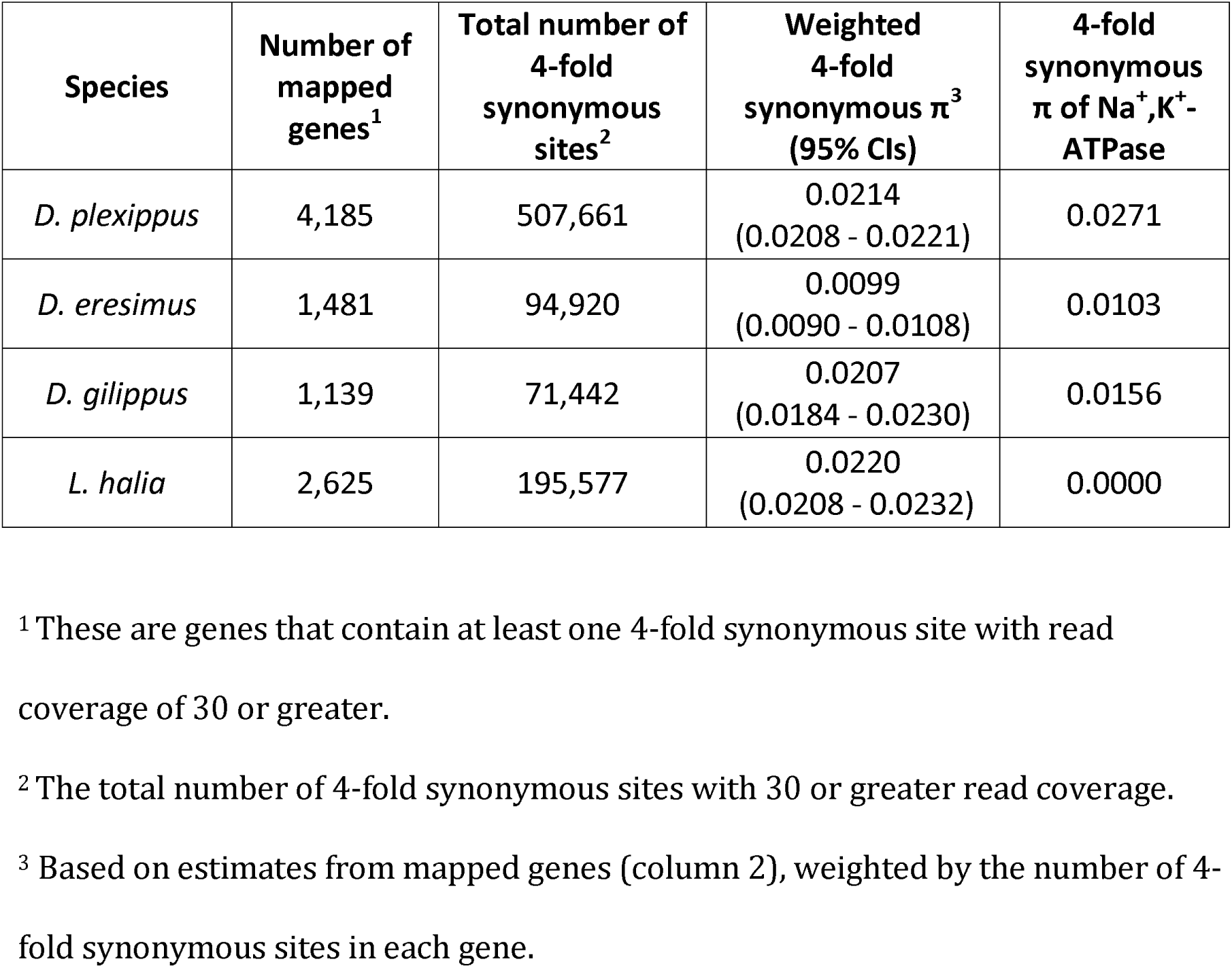
Levels of 4-fold synonymous nucleotide diversity (π_4f_) across all mapped 4-fold synonymous sites, as well as π_4f_ for Na^+^,K^+^-ATPase.

| Species | Number of mapped genes<sup>1</sup> | Total number of 4-fold synonymous sites<sup>2</sup> | Weighted 4-fold synonymous $\pi^3$ (95% CIs) | 4-fold synonymous $\pi$ of Na<sup>+</sup>,K<sup>+</sup>-ATPase |
| --- | --- | --- | --- | --- |
| *D. plexippus* | 4,185 | 507,661 | 0.0214<br>(0.0208 - 0.0221) | 0.0271 |
| *D. eresimus* | 1,481 | 94,920 | 0.0099<br>(0.0090 - 0.0108) | 0.0103 |
| *D. gilippus* | 1,139 | 71,442 | 0.0207<br>(0.0184 - 0.0230) | 0.0156 |
| *L. halia* | 2,625 | 195,577 | 0.0220<br>(0.0208 - 0.0232) | 0.0000 |

<sup>1</sup> These are genes that contain at least one 4-fold synonymous site with read coverage of 30 or greater.
<sup>2</sup> The total number of 4-fold synonymous sites with 30 or greater read coverage.
<sup>3</sup> Based on estimates from mapped genes (column 2), weighted by the number of 4-fold synonymous sites in each gene.

For Na^+^,K^+^-ATPase, the tree topology of this gene matched that observed from the full gene datasets (Fig. S2). However, species’ branch lengths for this gene were much shorter than in the full datasets and bootstrap support for this relationship was only 58%. Furthermore, this topology was not statistically better than either of the alternative topologies (Table 1). These observations indicate that this gene is highly conserved between these butterflies.

### Phylogenetic discordance

We carried out an analysis of gene tree discordance using BUCKy. This analysis revealed that the primary concordance tree was the same as that determined in the maximum-likelihood analyses using the reference-mapped and *de novo*-assembled concatenated, genomic datasets (Fig. 1D). However, the BUCKy analysis also revealed significant discordance (>5%) between individual gene trees and the inferred species tree. A higher proportion of genes supported a closer relationship between *D. gilippus* and *D. plexippus*, than between *D. gilippus* and *D. eresimus*.

The full dataset contained 5,735 informative sites for the ABBA/BABA test. The percentages of sites supporting each of the three possible relationships are given in Fig. 2. The D statistic for this data was ‐0.118 (95% CIs: ‐0.121 to ‐0.114). There were 4,869 4-fold synonymous sites in our reduced dataset. For this data, the D statistic was ‐0.134 (95% CIs: ‐0.138 to ‐0.130). There were 866 0-fold, non-synonymous sites. The D statistic was ‐0.025 (95% CIs: ‐0.031 to ‐0.019).

**Figure 2.**
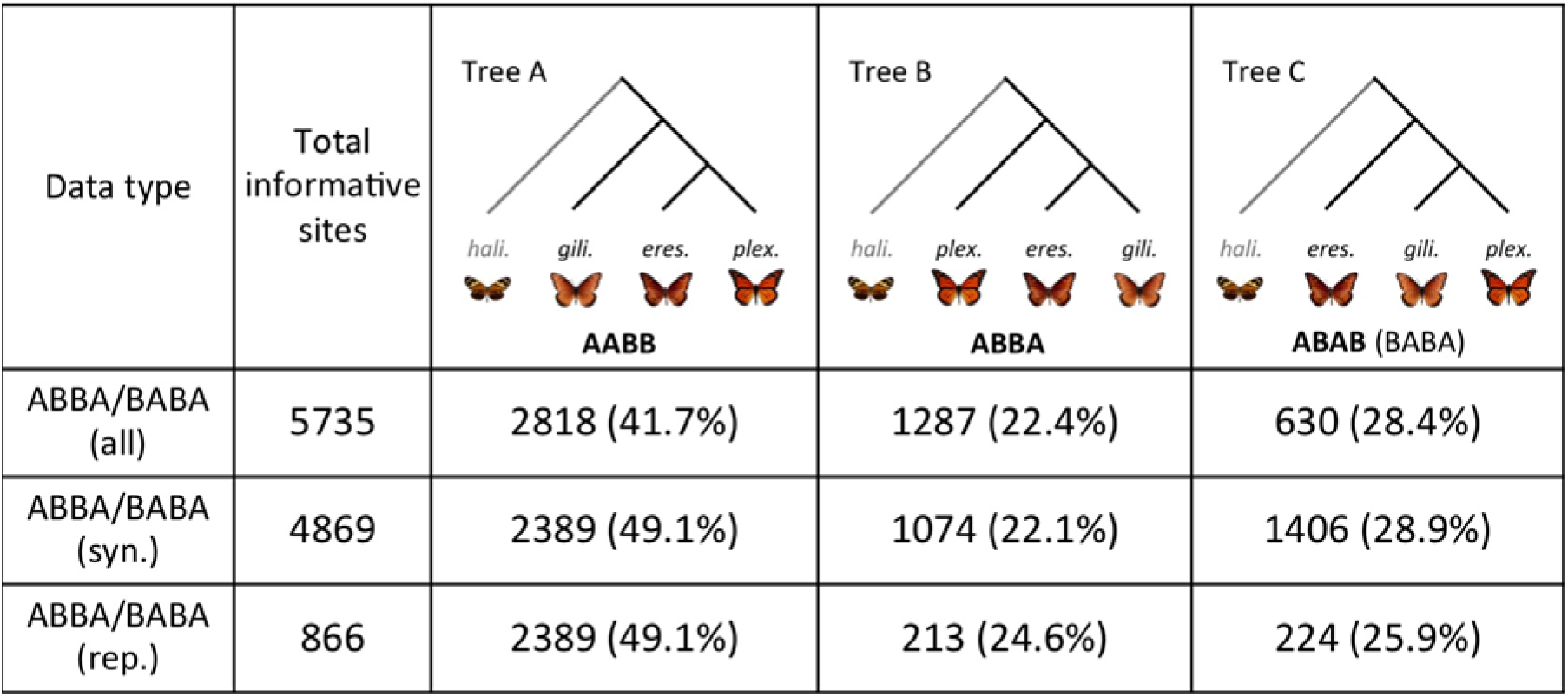
The percentages of ABBA/BABA informative sites that match the best-supported species tree (Tree A) and the two alternative trees for the three North American *Danaus* species in this study for either all 4-fold/0-fold sites (all), just 4-fold synonymous sites (syn.) or just 0-fold replacement sites (rep.), ('gili.' = *D. gilippus*, ‘eres.’ = *D. eresimus*, ‘plex’ = *D. plexippus*, ‘hali’ = *L. halia*).

For all classes of sites, the D statistic was significantly less than 0, indicating that ABBA and BABA sites were not equally represented. In all cases, this was due to a larger number of BABA sites than ABBA sites. This suggests that a greater amount of gene flow has occurred between *D. plexippus* and *D. gilippus* after splitting than between *D. eresimus* and *D. gilippus*. It is also possible that this result is due to structure in the ancestral populations of these species (Green et al. 2010).

### Genetic diversity and divergence time estimates

As rapid radiation increases the likelihood that ILS will occur between species (Whitfield and Lockhart 2007), we wanted to calculate the divergence times of these *Danaus* butterflies. However, a species’ effective population size (N_e_) can influence substitution rates (Ohta 1993; Woofit 2009), which could impact divergence time estimates. To assess the relative contemporary N_e_ of these species, we compared levels of 4-fold synonymous site diversity (π_4f_, Table 2). Of the four species, *D. eresimus* had the lowest synonymous diversity (π_4f_ = 0.010, 95% CIs: 0.009 – 0.011). The other three species all had similar levels of diversity (*L. halia*: π_4f_ = 0.022, 95% CIs: 0.021 – 0.023; *D. gilippus:* π_4f_ = 0.021, 95% CIs: 0.018 – 0.023; *D. plexippus*: π_4f_ = 0.021, 95% CIs: 0.021 – 0.022). As the *D. eresimus* dataset had fewer assembled transcripts (Table S2), we subsampled our *D. plexippus* dataset 10,000 times to calculate a mean and confidence intervals with the same number of genes as in the *D. eresimus* dataset. Our subsampled estimate of diversity for *D. plexippus* was significantly different from that of *D. eresimus* (π_4f_ = 0.021, 95% CIs: 0.020 – 0.022), indicating that the lower diversity level of *D. eresimus* is not likely due to the smaller number of genes represented in this dataset.

The similarities in genetic diversity levels exhibited in *D. plexippus* and *D. gilippus* suggest that they have comparable contemporary N_e_, while the population from which the *D. eresimus* sample was collected likely has a relatively smaller N_e_. Differences in N_e_ between these species could affect estimates of divergence times. Specifically, *D. eresimus* may have had a higher rate of evolution over time than the other three species. However, Tajima’s relative rate test (Tajima 1993), using all sites from the concatenated *de novo* dataset suggests that *D. eresimus* has had a similar rate of evolution to *D. gilippus* (χ^2^ = 2.46, p=0.074), whereas the rate of evolution in *D. plexippus* was higher than both *D. gilippus* (χ^2^ = 31.11, p<0.001) and *D. eresimus* (χ^2^ = 20.25, p<0.001). Thus, we do not infer any evolutionary patterns in relation to contemporary N_e_ in these species.

The estimate of pairwise synonymous divergence (dS) between *D. gilippus* and *D. eresimus* was 0.369 (95% CIs: 0.348 – 0.390). Assuming five generations per year (Malcolm et al. 1987), we estimated that the splitting time for *D. gilippus* and *D. eresimus* was 11.7 million years ago (MYA), (95% CIs: 11.0 – 12.4 MYA). For *D. gilippus* and *D. plexippus*, dS=0.312 (95% CIs: 0.310 – 0.337), and the estimated splitting time was 10.4 MYA (95% CIs: 9.7 – 10.8 MYA). The difference between these two divergence time estimates involving *D. gilippus* may reflect the larger extent of post-splitting genetic exchange that we infer to have occurred between *D. gilippus* and *D. plexippus*. It may also reflect a greater number of fixed differences driven by the relatively smaller N_e_ of *D. eresimus*. Synonymous divergence between *D. plexippus* and *D. eresimus* was 0.233 (95% CIs: 0.224 – 0.242), and our estimate of the splitting time between these species was 7.1 MYA (95% CIs: 6.8 – 7.4 MYA).

Bayesian estimates of divergence times using 4-fold synonymous sites and a log-normally distributed mutation rate suggested that *D. plexippus* and *D. eresimus* diverged 7.2 MYA (95% highest posterior density [HPD]: 6.9 – 7.5), whereas *D. gilippus* diverged from the common ancestor of *D. plexippus* and *D. eresimus* 11.0 MYA (95% HPD: 10.7 – 11.4). These estimates of splitting time are nearly identical to our simple estimates, which were also based on synonymous divergence and a fixed mutation rate. The simple divergence time estimate for *D. plexippus* and *D. eresimus* based on observed synonymous divergence falls within the 95% HPD interval for the divergence time between these species in the Bayesian analysis. The HPD interval for the divergence time of *D. gilippus* and the *D. plexippus/D. eresimus* common ancestor is somewhat lower than the simple estimate for the split between *D. gilippus* and *D. eresimus*, but somewhat higher than the estimate for that between *D. gilippus* and *D. plexippus*. Overall, these divergence estimates suggest that the three species diverged relatively rapidly from one another (within 3-4 million years), which may have increased the likelihood that ILS occurred.

We explicitly examined the extent of ILS given our estimated divergence times by simulating the coalescent of 10,000 genes in *ms* (Hudson 2002). We used 0.02 for our estimate of θ (4N_e_μ) and converted our estimated divergence times to units of 4N_e_generations (run command: ms 3 10000 ‐t 0.02 ‐I 3 1 1 1 ‐ej 2.4 1 2 ‐ej 3.6 2 3 ‐T). From these simulations, we infer that ~8% of genetic regions will exhibit an alternative topology to that of the true species relationship given the divergence times calculated here.

## Discussion

Evolutionary relationships within the *Danaus* genus are a subject of uncertainty due to incongruences in both morphology and genetic sequences (Ackery and Vane-Wright 1984; Kitching 1985; Brown et al. 2004; Smith et al. 2005; Zhen et al. 2012). In contrast to previous studies (Ackery and Vane-Wright 1984; Lushai et al. 2003; Smith et al. 2005; Brower et al. 2010), the analyses described here indicate that *D. plexippus* and *D. eresimus* are the more closely related species, whereas *D. gilippus* is a more distant relative. While this relationship is well supported, we also observe significant phylogenetic discordance between these species. This may be attributed to several factors including incomplete lineage sorting (ILS) and post-speciation gene flow. Relative to the amount of evolutionary divergence that occurred after these three species split from their common ancestor, their speciation times are brief. This means that ILS is likely to explain some phylogenetic incongruence. Using coalescence simulations, we calculated that ~8% of gene regions will exhibit an alternative topology to that of the true species tree given our estimated splitting times. This is less than the ~18% of phylogenetic discordance detected. This observation, combined with the results of the ABBA/BABA test, suggests that genetic exchange after these species diverged also likely contributed to incongruences between these species. Similar explanations have been proposed to explain extensive phylogenetic discordance in many other species groups (e.g. Moody and Rieseberg 2012; Cui et al. 2013; Liu et al 2015), including butterflies (Kozak et al. 2015).

In addition to amending current phylogenetic understanding of the *Danaus* genus, our results shed light on the origins of the N122H substitution in the H1-H2 extracellular domain of Na^+^,K^+^-ATPase. As the gene tree topology for this region is concordant with the best-supported phylogeny from whole nuclear gene datasets (Fig. S2), the most parsimonious scenario to explain the N122H mutation in both *D. plexippus* and *D. eresimus* is an emergence in their shared common ancestor. If this is the case, it then either fixed before their split, or else fixed independently in each lineage (Fig. S3A). It is also possible that the N122H mutation arose in the common ancestor of all three *Danaus* species, and was subsequently lost in *D. gilippus* (Fig. S3B). We cannot confidently differentiate between this and the previous hypothesis with the data currently available.

If the N122H allele had a strong adaptive advantage when it arose, it most likely fixed rapidly. This would have resulted in a selective sweep, reducing genetic diversity in this gene (Maynard Smith and Haigh 1974). Interestingly, 4-fold synonymous diversity estimates for Na^+^,K^+^-ATPase in both *D. plexippus* and *D. eresimus* are slightly higher than genome-wide estimates (Table 2). This suggests that if a sweep occurred, it was not recent in either species’ evolutionary history.

Estimated synonymous divergence (dS) between *D. plexippus* and *D. eresimus* for Na^+^,K^+^-ATPase is 0.15, whereas the genome-wide estimate is 0.23. This divergence estimate translates to a splitting time of ~5.4 MY for Na^+^,K^+^-ATPase, which is less than the estimated splitting time for these two species based on the full gene dataset. This observation lends support to the third hypothesis that adaptive introgression from either *D. plexippus* or *D. eresimus* into the other species can explain the presence of the N122H mutation in both butterflies (Fig. S3C). However, had this occurred, we would predict that Na^+^K^+^-ATPase genetic diversity in one of these species would be low relative to other genetic regions. As noted previous, we actually observe relatively high levels of synonymous diversity in Na^+^K^+^-ATPase. Additionally, in the case of adaptive introgression, divergence in this gene between *D. plexippus* and *D. eresimus* should be low relative to other genes. While we do observe this (Fig. S2, Fig. S4), Na^+^K^+^-ATPase divergence is low overall among all the butterflies in this study. This suggests that the low level of Na^+^K^+^-ATPase divergence between *D. plexippus* and *D. eresimus* is not the result of introgression, but rather that this gene is highly conserved between these butterflies.

Another question this study raises is the origins of the extensive phenotypic divergence observed in *D. plexippus*. Divergent traits of *D. plexippus* include changes in wing size, color and mating strategy, as well as the establishment of long-distance migration (Ackery and Vane-Wright 1984). Changes in wing coloration may have arisen in response to selection for increased mimic resemblance with *Limenitis archippus* (the viceroy), as these two butterflies are Müllerian mimics of one another (Ritland and Brower 1991). Other behavioral and morphological shifts could have occurred in response to the evolution of migratory behavior in *D. plexippus* (Wells et al. 1993; Dockx 2007). It is interesting in this respect that recent work suggests all contemporary monarch populations (including non-migratory ones) originated from a North American migratory population (Brower et al. 2007; Zhan et al. 2014).

Changes in mating strategy, which could have arisen in connection with migratory behavior, may also have produced some morphological changes (Ackery and Vane-Wright 1984). The uniquely derived traits of *D. plexippus* have likely obscured phylogenetic relationships between North American *Danaus* species and lead to the different conclusions than the ones presented here. It is noteworthy in this regard that Tajima’s relative rate test (based on all sites) suggests that *D. plexippus* has experienced a significantly higher rate of molecular evolution than the other two *Danaus* species (see Results).

It is also of interest that in most previous genetic analyses of *Danaus* species relationships, all *D. eresimus* samples and nearly all *D. gilippus* samples came from the Cayman Islands (Lushai et al. 2003; Smith et al. 2005; Brower et al. 2010). It is possible that these samples may have unique attributes (e.g. recent genetic exchange, reduced N_e_, etc.), and are not typical of their species more broadly. While the samples in this study all came from continental North America, they may also not accurately capture the genetic diversity patterns of these widespread butterflies. More specifically, it is possible that either contemporary or ancestral population structure could be confounding some of our results. Population structure within species can generate genetic patterns that resemble those of hybridization between species (Green et al. 2010; Eriksson and Manica 2012). Very little population structure has been observed in *D. plexippus*, presumably due to their migratory behavior and overall high mobility (Eanes and Koehn 1978, Brower and Boyce 1991, Lyons et al. 2012). Less is known about population structure in *D. gilippus* or *D. eresimus*. However, while not migratory like *D. plexippus*, both are also considered to be highly mobile (Smith et al. 2002). In future work, we hope to address interesting questions regarding gene flow and population structure within these butterflies throughout their geographic ranges. We also hope to look across the *Danaus* clade more broadly to assess levels of phylogenetic incongruence throughout this genus.

## Conclusions

Common ancestry is the simplest explanation for the shared N122H mutation in the H1-H2 extracellular domain of Na^+^,K^+^-ATPase observed in *D. plexippus* and *D. eresimus*. However, earlier origins within *Danaus* or post-speciation genetic exchange are also possibilities. Relatively rapid divergence and historical introgression likely contributed to observed phylogenetic incongruences among these North American *Danaus* butterflies. As the majority of phylogenetic analyses for the milkweed butterflies (*Danaidae*) have relied heavily on morphology and limited genetic data, this study highlights a potential need to reevaluate the relationships among milkweed butterflies more broadly. In particular, the extent of incomplete lineage sorting and genetic introgression in this clade should be assessed. Such examinations could potentially reveal more about the evolution of novel morphological and behavioral traits in these butterflies including *D. plexippus*.

## Acknowledgements

V. Miró Pina and M. Schumer provided helpful comments on earlier versions of this work. A Briscoe kindly contributed the *D. gilippus, D. eresimus* and *L. halia* for the original study that produced the original data utilized here.

## Data Accessibility

The data used for this study can be found at: http://genomics-pubs.princeton.edu/insect_genomics/data.shtml

## Supplementary Information

**Table S1.**
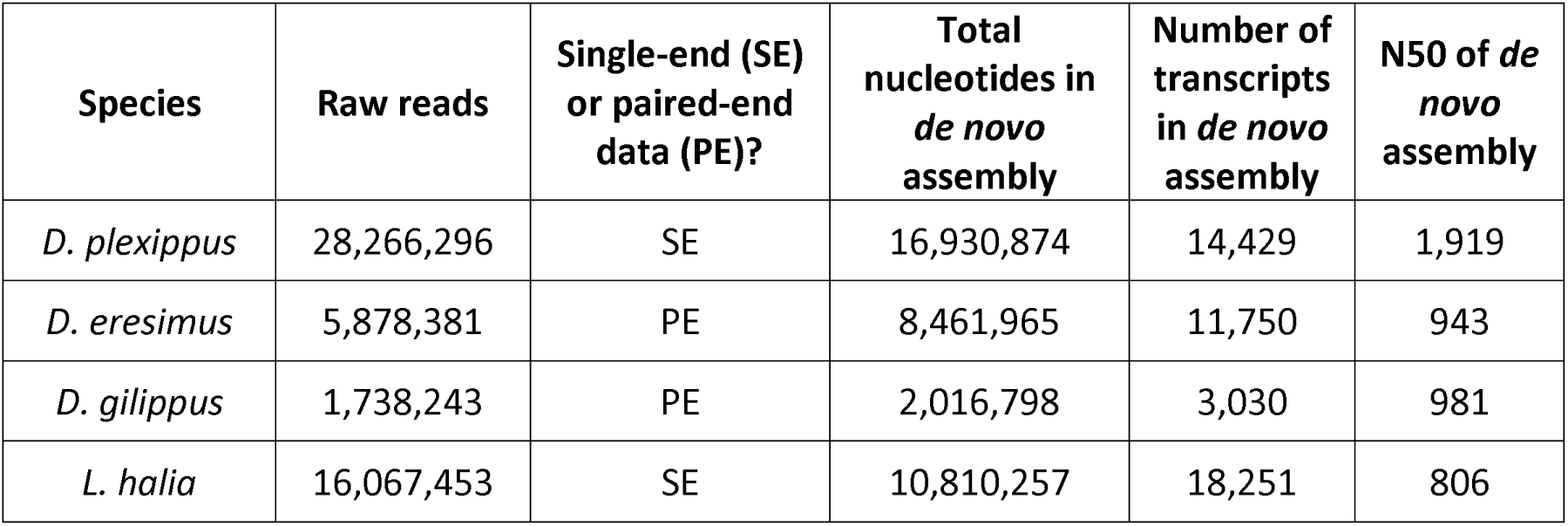
Summary statistics for the raw data used in this study, as well as the *de novo* assembly of each species’ transcriptome.

| Species | Raw reads | Single-end (SE) or paired-end data (PE)? | Total nucleotides in *de novo* assembly | Number of transcripts in *de novo* assembly | N50 of *de novo* assembly |
| --- | --- | --- | --- | --- | --- |
| *D. plexippus* | 28,266,296 | SE | 16,930,874 | 14,429 | 1,919 |
| *D. eresimus* | 5,878,381 | PE | 8,461,965 | 11,750 | 943 |
| *D. gilippus* | 1,738,243 | PE | 2,016,798 | 3,030 | 981 |
| *L. halia* | 16,067,453 | SE | 10,810,257 | 18,251 | 806 |

**Table S2.** The total number of Illumina reads for each species after quality trimming, plus the number or trimmed reads that mapped to either the *D. plexippus* CDS or a species-specific, *de novo* assembled transcriptome.

| Species | Total trimmed reads | Reads mapped to *D. plexippus* reference CDS |  | Reads mapped to *de novo* assembled transcriptome |  |
| --- | --- | --- | --- | --- | --- |
| *D. plexippus* | 27,068,642 | mapped | 19,132,116 | mapped | 25,598,815 |
|  |  | percent | 70.7% | percent | 94.6% |
| *D. eresimus* | 5,354,186 (pairs) | mapped | 2,021,741 | mapped | 4,324,576 |
|  |  | percent | 37.9% | percent | 80.8% |
| *D. gilippus* | 1,538,115 (pairs) | mapped | 1,268,745 | mapped | 1,523,195 |
|  |  | percent | 82.5% | percent | 99.1% |
| *L. halia* | 14,530,186 | mapped | 11,649,577 | mapped | 14,075,972 |
|  |  | percent | 80.2% | percent | 96.9% |

**Table S3.** Percentage of gene trees that support each of the three alternative tree topologies possible among the *Danaus* species in this study using an a prior of either 0.1,1.0 or 10. Concordance factors (CF) are presented with creditability intervals (Cl) in parentheses.

| Species Relationships | CF with $\alpha = 0.1$ (CI) | CF with $\alpha = 1.0$ (CI) | CF with $\alpha = 10.0$ (CI) |
| --- | --- | --- | --- |
| Tree A (Fig. 2) | 0.824 (0.772 - 0.874) | 0.822 (0.770 - 0.872) | 0.804 (0.753 - 0.851) |
| Tree B (Fig. 2) | 0.057 (0.023 - 0.092) | 0.058 (0.025 - 0.094) | 0.069 (0.038 - 0.103) |
| Tree C (Fig. 2) | 0.119 (0.070 - 0.165) | 0.120 (0.075 - 0.165) | 0.127 (0.086 - 0.172) |

## Supplementary Figures

**Figure S1.**
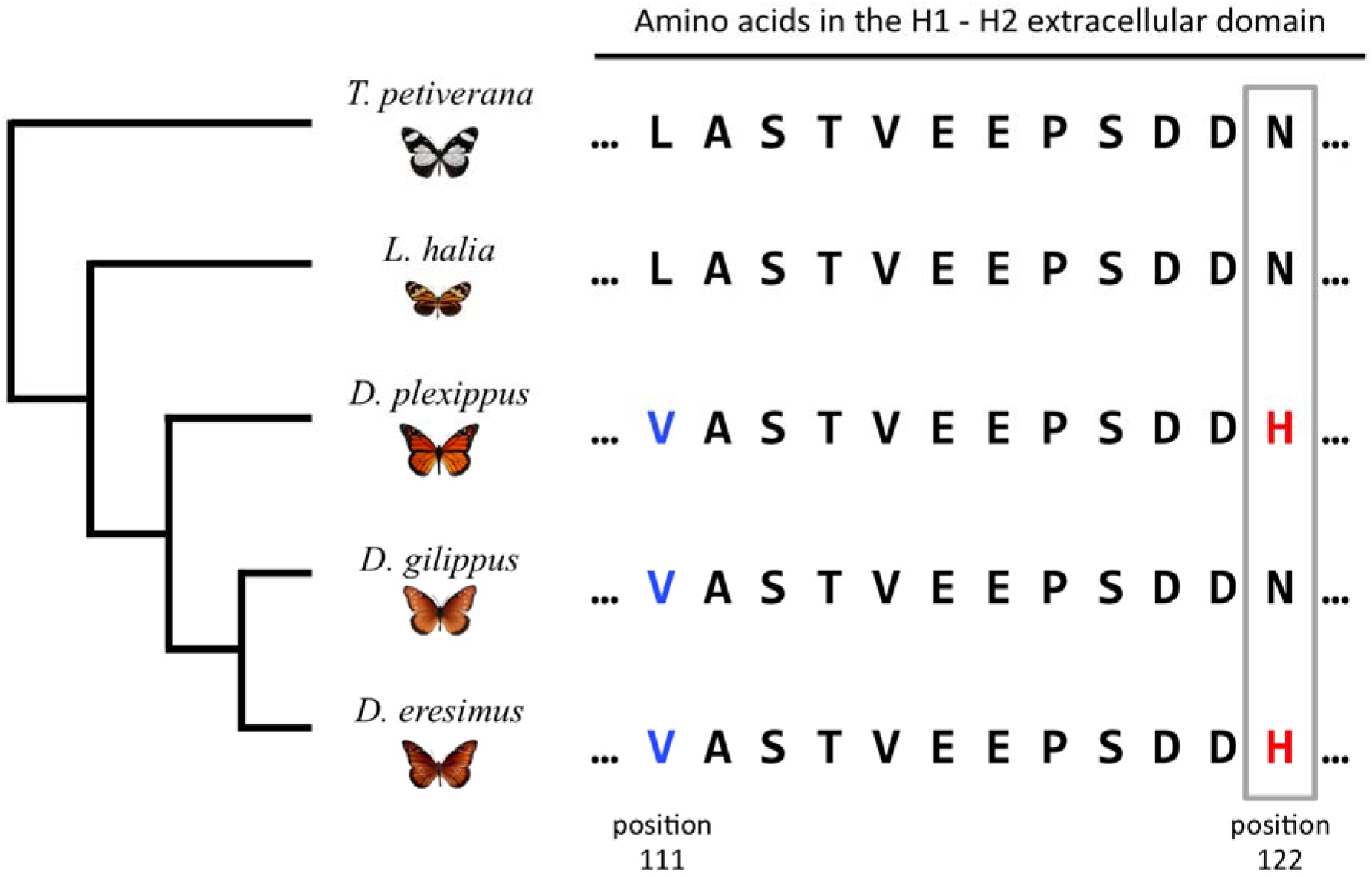
Historical phylogeny (see Ackery and Vane-Wright 1984; Lushai et al. 2003; Smith et al. 2005; Brower et al. 2010) and amino acid sequences of the H1-H2 extracellular domain of the Na^+^,K^+^-ATPase gene in the *Danaus* species examined in this study, plus a second milkweed butterfly outgroup *Tirumala petiverana* (adapted from Aardema et al. 2012, Zhen et al. 2012). The presence of a Valine at position 111 (blue letters) in the three *Danaus* butterflies likely facilitates their ability to feed on milkweed and possibly sequester cardenolides. The N122H mutation (red letters) seen in *D. plexippus* and *D. eresimus* may also contribute to cardenolide feeding and sequestration ability (Zhen et al. 2012; Petschenka et al. 2013).

**Figure S2.**
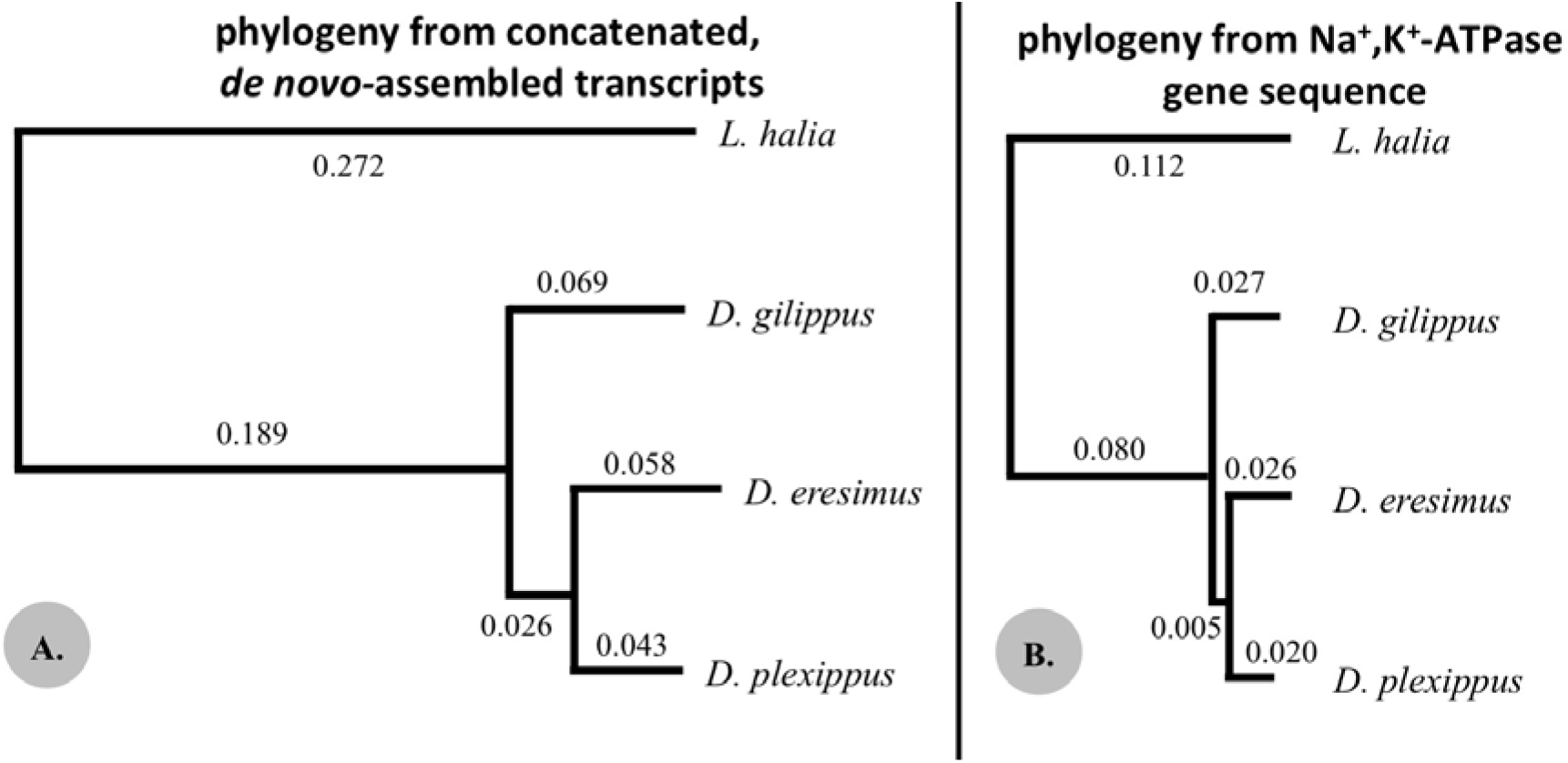
A.) The best supported phylogenetic relationship as determined using data from 478 *de novo*-assembled nuclear gene regions. Numbers indicate branch lengths (this is the same tree as shown in Fig. 1C). B.) The best supported phylogenetic relationship produced with data from the Na^+^,K^+^-ATPase gene (in RAxML with a gamma distribution of rate heterogeneity [i.e. GTR + *T* model]). This relationship has 58% bootstrap support based on 100 bootstrap replicates. The branches of these two trees are drawn to the same scale.

**Figure S3.**
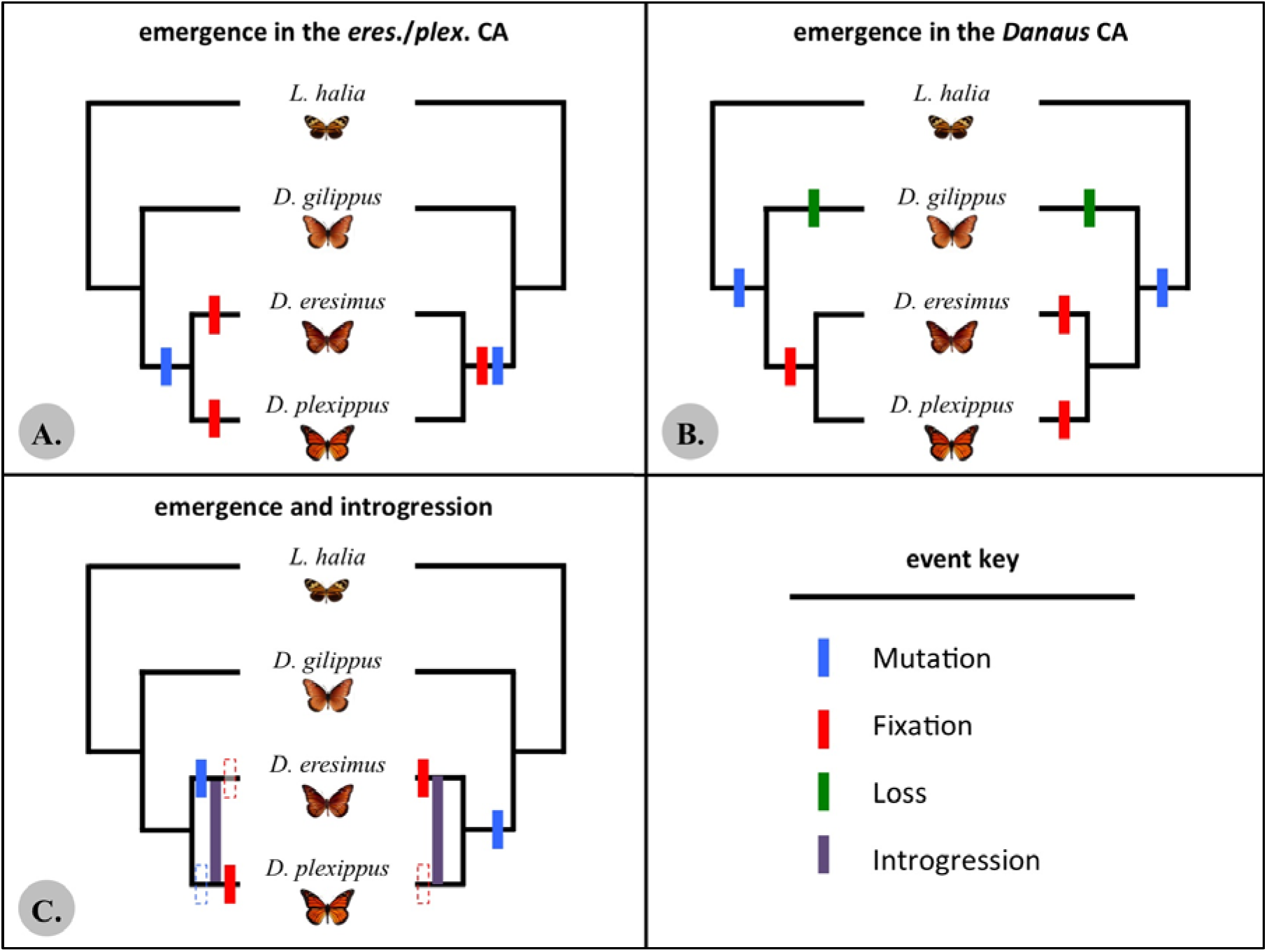
Graphical representation of potential scenarios to account for the presence of the N122H mutation in the H1-H2 extracellular domain of Na^+^,K^+^‐ ATPase in both *D. plexippus* and *D. eresimus*. Not all possible scenarios are displayed in this figure. A) The N122H mutation arose in the common ancestor of *D. plexippus* and *D. eresimus* and either fixed in this common ancestor (right), or else fixed independently in each linage (left). B) The N122H mutation arose in the common ancestor of all three species, but was lost in *D. gilippus*. C) The mutation arose or was present in either *D. plexippus* or *D. eresimus*, and then introgressed into the second species. The dashed boxes are present to indicate that we cannot infer the direction of this potential introgression, although our data suggests that if introgression is responsible for this mutation in both species, then it is more probably that it introgressed from *D. plexippus* into *D. eresimus*.

**Figure S4.**
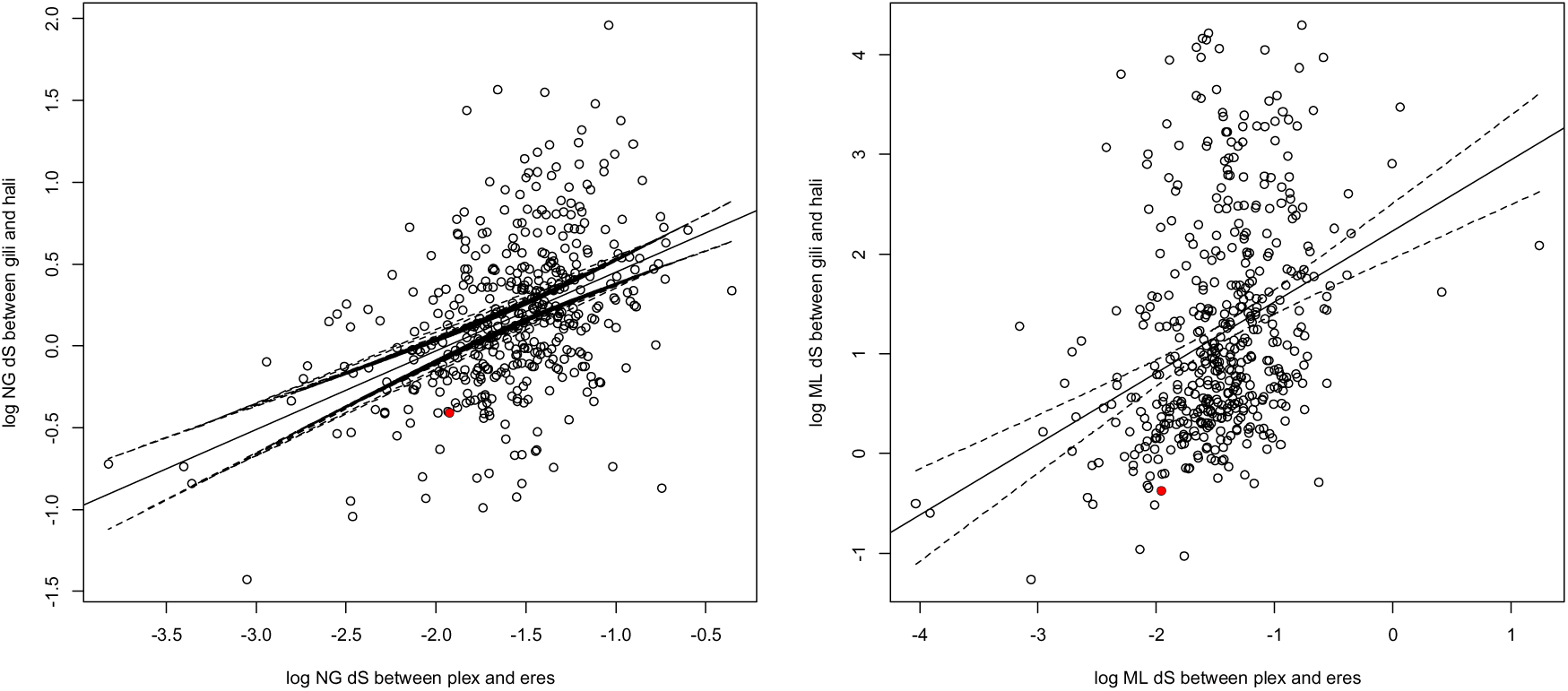
Comparison of log dS between *D. plexippus* and *D. eresimus* (x axis), and log dS between *D. gilippus* and *L. halia* (y axis). The left figure shows dS estimates based on the method of Nei and Gojobori (1986), as determined in PAML (v. 4.8; Yang 2007). The right figure shows maximum-likelihood estimates of dS, also determined in PAML. The single red dot in each figure indicates the location of the dS estimates for the Na^+^,K^+^-ATPase gene. Solid lines indicate the regression line for each dataset, and the dashed lines are the 95% confidence intervals for this line.

## References

Aardema, M. L., Y. Zhen and P. Andolfatto. 2012. The evolution of cardenolide-resistant forms of Na^+^,K^+^-ATPase in Danainae butterflies. Mol. Ecol. 21: 340–349.

Ackery, P. R. and R. I. Vane-Wright. 1984. Milkweed butterflies: their cladistics and biology. Cornell Univ. Press, Ithaca, NY.

Agrawal, A. A., G. Petschnka, R. A. Bingham, M. G. Weber and S. Rasmann. 2012. Toxic cardenolides: chemical ecology and coevolution of specialized plant–herbivore interactions. New Phytol 194:28–45.

Ané, C., B. Larget, D. A. Baum, S. D. Smith and A. Rokas. 2007. Bayesian estimation of concordance among gene trees. Mol. Biol. Evol. 24:412–426.

Altschul, S. F., W. Gish, W. Miller, E. W. Myers and D. J. Lipman. 1990. Basic local alignment search tool. J. Mol. Biol. 215:403–410.

Baack, E. J. and L. H. Rieseberg. 2007. A genomic view of introgression and hybrid speciation. Curr. Opin. Genet. Dev. 17:513–518.

Bachtrog, D., K. Thornton, A. Clark and P. Andolfatto. 2006. Extensive introgression of mitochondrial DNA relative to nuclear genes in the Drosophila yakuba species group. Evolution 60:292–302.

Brower, A. V. Z. and T. M. Boyce. 1991. Mitochondrial DNA variation in monarch butterflies. Evolution 45:1281–1286.

Brower, L. P. and C. M. Moffit. 1974. Palatability dynamics of cardenolides in the monarch butterfly. Nature 249:280–283.

Brower, L. P., K. S. Oberhauser, M. Boppré, A. V. Z. Brower and R. I. Vane-Wright. 2007. Monarch sex: ancient rites, or recent wrongs? Antenna 31:12–18.

Brower, A. V. Z., N. Wahlberg, J. R. Ogawa, M. Boppré, and R. I. Vane-Wright. 2010. Phylogenetic relationships among genera of danaine butterflies (Lepidoptera: Nymphalidae) as implied by morphology and DNA sequences. Syst. Biodivers. 8:75–89.

Brown, K. S. Jr., B. von Schoultz and E. Suomalainen. 2004. Chromosome evolution in Neotropical Danainae and Ithomiinae (Lepidoptera). Hereditas 141:216–236.

Charlesworth, B. 2009. Effective population size and patterns of molecular evolution and variation. Nat. Rev. Genet. 10:195–205

Cui, R., M. Schumer, K. Kruesi, R. Walter, P. Andolfatto and G. G. Rosenthal. 2013. Phylogenomics reveals extensive reticulate evolution in Xiphophorus fishes. Evolution 67:2166–2179.

Darriba, D., G. L. Taboada, R. Doallo and D. Posada. 2012. jModelTest 2: more models, new heuristics and parallel computing. Nat. Methods 9:772.

Degnan, J. H. and N. A. Rosenberg. 2006. Discordance of species trees with their most likely gene trees. PLoS Genet. 2:e68.

Degnan, J. H. and N. A. Rosenberg. 2009. Gene tree discordance, phylogenetic inference and the multispecies coalescent. Trends Ecol. Evol. 24:332–340.

Dobler, S., G. Petschenka and H. C. Pankoke. 2011. Coping with toxic plant compounds – the insect's perspective on iridoid glycosides and cardenolides. Phytochemistry 72:1593–1604.

Dobler, S., S. Dalla, V. Wagschal and A. A. Agrawal. 2012. Community-wide convergent evolution in insect adaptation to toxic cardenolides by substitutions in the Na,K-ATPase. Proc. Natl. Acad. Sci. USA 109:13040–13045.

Dobler, S., G. Petschnka, V. Wagschal, L. Flacht. 2015. Convergent adaptive evolution – how insects master the challenge of cardiac glycoside-containing host plants. Entomol. Exp. Appl. 157:30–39.

Dockx, C. 2007. Directional and stabilizing selection on wing size and shape in migrant and resident monarch butterflies, Danaus plexippus (L.), in Cuba. Biol. J. Linn. Soc. 92:605–616.

Drummond, A. J. and A. Rambaut. 2007. BEAST: Bayesian evolutionary analysis by sampling trees. BMC Evol. Biol. 7:214.

Eanes, W.F. and R. K. Koehn. 1978. Analysis of genetic structure in the monarch butterfly, Danaus plexippus L. Evolution 32:784–797.

Edgar, R. C. 2004. MUSCLE: multiple sequence alignment with high accuracy and high throughput. Nucleic Acids Res. 32:1792–7197.

Eriksson, A. and A. Manica. 2012. Effect of population structure on the degree of polymorphism shared between modern human populations and ancient hominins. Proc. Natl. Acad. Sci. USA 109:13956–13960.

Gernhard, T. 2008. The conditioned reconstructed process. J. Theor. Biol. 253:769–778.

Gouy, M., S. Guindon and O.. Gascuel. 2010. SeaView version 4: a multiplatform graphical user interface for sequence alignment and phylogenetic tree building. Mol. Biol. Evol. 27:221–224.

Green, R. E., J. Krause, A. W. Briggs, T. Maricic, U. Stenzel, M. Kircher, N. Patterson, H. Li, W. Zhai, M. H. Fritz, et al. 2010. A draft sequence of the Neanderthal genome. Science 328:710–722.

Guindon, S. and O. Gascuel. 2003. A simple, fast and accurate method to estimate large phylogenies by maximum-likelihood. Syst. Biol. 52:696–704.

Hasegawa, M., H. Kishino and T. Yano. 1985. Dating of the human-ape splitting by a molecular clock of mitochondrial DNA. J. Mol. Evol. 22:160–174.

Hershberg, R. and D. A. Petrov. 2008. Selection on codon bias. Annu. Rev. Genet. 42:287–299.

Holzinger, F. and M. Wink. 1996. Mediation of cardiac glycoside insensitivity in the monarch butterfly (Danaus plexippus): role of an amino acid substitution in the ouabain binding site of Na^+^,K^+^-ATPase. J. Chem. Ecol. 22:1921–1937.

Hornett, E. A. and C. W. Wheat. 2012. Quantitative RNA-Seq analysis in non-model species: assessing transcriptome assemblies as a scaffold and the utility of evolutionary divergent genomic reference species. BMC Genomics 13:361.

Hudson, R. R. 2002. Generating samples under a Wright-Fisher neutral model of genetic variation. Bioinformatics 18: 337–338.

Hudson, R. R., M. Kreitman and M. Aguadé. 1987. A test of neutral molecular evolution based on nucleotide data. Genetics 116:153–159.

Huelsenbeck, J. P., B. Larget and M. E. Alfaro. 2004. Bayesian phylogenetic model selection using reversible jump Markov chain Monte Carlo. Mol. Biol. Evol. 21:1123–1133.

Keightley, P. D., A. Pinharanda, R. W. Ness, F. Simpson, K. K. Dasmahapatra, J. Mallet, J. W. Davey and C. D. Jiggins. 2015. Estimation of the spontaneous mutation rate in Heliconius melpomene. Mol. Biol. Evol. 32:239–243.

Kozak, K. M., N. Wahlberg, A. Neild, K. K. Dasmahapatra, J. Mallet and C. D. Jiggins. 2015. Multilocus species trees show the recent adaptive radiation of the mimetic Heliconius butterflies. Syst. Bio. 64:505–524.

Kunsch, H. 1989. The jackknife and the bootstrap for general stationary observations. Ann. Stat. 17:1217–1241.

Larget, B. R., S. K. Kotha, C. N. Dewey and C. Ané. 2010. BUCKy: Gene tree/species tree reconciliation with Bayesian concordance analysis. Bioinformatics 26:2910–2911.

Lawrie, D. S., P. W. Messer, R. Hershberg and D. A. Petrov. 2013. Strong purifying selection at synonymous sites in D. melanogaster. PLoS Genet. 9:e1003527.

Li, H., B. Handsaker, A. Wysoker, T. Fennell, J. Ruan, N. Homer, G. Marth, G. Abecasis, R. Durbin and 1000 Genome Project Data Processing Subgroup. 2009. The Sequence alignment/map (SAM) format and SAMtools. Bioinformatics 25:2078–2079

Liu, K. J., E. Steinberg, A. Yozzo, Y. Song, M. H. Kohn and L. Nakhleh. 2015. Interspecific introgressive origin of genomic diversity in the house mouse. Proc. Natl. Acad. Sci. USA 112:196–201.

Lunter, G. and M. Goodson. 2011. Stampy: a statistical algorithm for sensitive and fast mapping of Illumina sequence reads. Genome Res. 21:936–939.

Lushai, G., D. A. S. Smith, D. Goulson, J. A. Allen and N. Maclean. 2003. Mitochondrial DNA clocks and the phylogeny of Danaus butterflies. Insect Sci. Appl. 23:309–315.

Lyons, J. I., A. A. Pierce, S. M. Barribeau, E. D. Sternberg, A. J. Mongue and J. C. de Roode. 2012. Lack of genetic differentiation between monarch butterflies with divergent migration destinations. Mol. Ecol. 21:3433–3444.

Maddison, W. P. and L. L. Knowles. 2006. Inferring phylogeny despite incomplete lineage sorting. Syst. Biol. 55:21–30.

Malcolm, S. B., B. J. Cockrell and L. P. Brower. 1987. Monarch butterfly voltinism: effects of temperature constraints at different latitudes. Oikos 49:77–82.

Mallet, J., N. Besansky and M. W. Hahn. 2015. How reticulated are species? Bioessays 37: DOI 10.1002/bies.201500149.

Maynard Smith, J. and J. Haigh. 1974. The hitchhiking effect of a favourable gene. Genet. Res. 23:23–35.

McKenna, A., M. Hanna, E. Banks, A. Sivachenko, K. Cibulskis, A. Kernytsky, K. Garimella, D. Altshuler, S. Gabriel, M. Daly and M. A. DePristo. 2010. The Genome Analysis Toolkit: a MapReduce framework for analyzing next-generation DNA sequencing data. Genome Res. 20:1297–1303.

Moody, M. L. and L. H. Rieseberg. 2012. Sorting through the chaff, nDNA gene trees for phylogenetic inference and hybrid identification of annual sunflowers [*Helianthus sect. Helianthus*). Mol. Phylogenet. Evol. 64:145–155.

Nei, M. 1987. Molecular evolutionary genetics. New York: Columbia Univ. Press.

Nei, M. and W-H. Li. 1979. Mathematical model for studying genetic variation in terms of restriction endonucleases, Proc. Natl. Acad. Sci. USA 76:5269–5273.

Nei, M. and T. Gojobori. 1986. Simple methods for estimating the numbers of synonymous and nonsynonymous nucleotide substitutions. Mol. Biol. Evol. 3:418–426.

Obbard, D. J., J. Maclennan, K-W. Kim, A. Rambaut, P. M. O’Grady and F. M. Jiggins. 2012. Estimating divergence dates and substitution rates in the *Drosophila* phylogeny. Mol. Biol. Evol. 29:3459–3473.

Ohta, T. 1993. Amino acid substitution at the Adh locus of Drosophila is facilitated by small population size. Proc. Natl. Acad. Sci. USA. 90:4548–51.

Petschenka, G., S. Fandrich, N. Sander, V. Wagschal, M. Boppré and S. Dobler. 2013. Stepwise evolution of resistance to toxic cardenolides via genetic substitutions in the Na+/K+-ATPase of milkweed butterflies (Lepidoptera: Danaini). Evolution 67:2753–2761.

Rambaut, A., M. A. Suchard, D. Xie and A. J. Drummond. 2014. Tracer vl.6, Available from http://beast.bio.ed.ac.uk/Tracer.

Ritland, D. B. and L. P. Brower. 1991. The viceroy butterfly is not a batesian mimic. Nature 350:497–498.

Ronquist, F., and J. P. Huelsenbeck. 2003. MrBayes 3: Bayesian phylogenetic inference under mixed models. Bioinformatics 19:1572–1574.

Schulz, M. H., D. R. Zerbino, M. Vingron and E. Birney. 2012. Oases: Robust de novo RNA-seq assembly across the dynamic range of expression levels. Bioinformatics 28:1086–1092.

Schwartz, R. S. and R. L. Mueller. 2010. Branch length estimation and divergence dating: estimates of error in Bayesian and maximum likelihood frameworks. BMC Evol. Biol. 10:5.

Shimodaira, H. and M. Hasegawa. 2001. CONSEL: for assessing the confidence of phylogenetic tree selection. Bioinformatics 17:1246–1247.

Smith, D. A. S., G. Lushai and J. A. Allen. 2005. A classification of *Danaus* butterflies (Lepidoptera: Nymphalidae) based upon data from morphology and DNA. Zool. J. Linn. Soc. 144:191–212.

Stamatakis, A. 2014. RAxML Version 8: A tool for Phylogenetic Analysis and Post-Analysis of Large Phylogenies. Bioinformatics 30:1312–1313.

Stern, D. L. 2013. The genetic causes of convergent evolution. Nat. Rev. Genet. 14:751–764.

Tajima, F. 1993. Simple methods for testing the molecular evolutionary clock hypothesis. Genetics 135:599–607.

Tavare, S. 1986. Some Probabilistic and Statistical Problems in the Analysis of DNA Sequences. Lectures Math. Life Sci. 17:57–86.

Wells, H., P. H. Wells and S. H. Rogers. 1993. Is multiple mating an adaptive feature of monarch butterfly winter aggregation? Pp. 61–68.

S. B. Malcolm and M. P. Zalucki, eds. Biology and Conservation of the Monarch Butterfly. No. 38 Science Series, Nat. Hist. Mus. of Los Angeles Co. Los Angeles, CA. USA.

Woofit, M. 2009. Effective population size and the rate and pattern of nucleotide substitutions. Biol. Lett. 5:417–420.

Yang, Z. 2007. PAML 4: a program package for phylogenetic analysis by maximum likelihood. Mol. Biol. Evol. 24: 1586–1591.

Zerbino, D.R. and E. Birney. 2008. Velvet: algorithms for de novo short read assembly using de Bruijn graphs. Genome Res. 18:821–829.

Zhan, S., W. Zhang, K. Niitepõld, J. Hsu, J. F. Haeger, M. P. Zalucki, S. Altizer, J. C. de Roode, S. M. Reppert and M. R. Kronforst. 2014. The genetics of monarch butterfly migration and warning coloration. Nature 514:317–321.

Zhen, Y., M. L. Aardema, E. M. Medina, M. Schumer and P. Andolfatto. 2012. Parallel molecular evolution in an herbivore community. Science 337:1634–1637.

